# Enrich.AI: An Agentic AI Platform for Interactive Gene Set Enrichment Analysis and Biological Interpretation

**DOI:** 10.64898/2026.09.29.755561

**Authors:** Dimitrios E. Koumadorakis, Themis P. Exarchos, Panagiotis Vlamos, Aristidis G. Vrahatis

## Abstract

**Motivation:** Enrichment analysis is widely used to extract biological meanings from gene lists. However, the large number of available tools and reference libraries often make the interpretation of results challenging. Inexperienced users lack clear guidance on which datasets and terms are most relevant to their data, whereas experienced researchers must navigate large result sets and consult multiple resources to identify biologically relevant findings. Integrating AI into this process can help to bridge these gaps.

**Results:** We present Enrich.AI, a conversational biology assistant that integrates statistical enrichment analysis with automated biological interpretation. The agent operates on an LLM-based ReAct architecture and dynamically selects among the tools for enrichment (220+ gene set libraries), literature search (40+ million publications via Europe PMC), gene annotation, and database retrieval. Given a gene list and a short description of the biological context, Enrich.AI produces a written interpretation grounded in the retrieved literature and six interactive, configurable visualizations.A conversational interface enables human-in-the-loop refinement, letting researchers without programming experience explore results iteratively. The reliability of these interpretations was validated through a systematic evaluation on 50 gene sets, demonstrating high faithfulness to retrieved evidence and a low hallucination rate.

**Availability and implementation:** Enrich.AI is freely available as a Streamlit web application at https://enrichai.streamlit.app with source code released at https://github.com/jkouprey/Enrich.AI.

**Supplementary information:** **Supplementary data are available online**.

## 1. Introduction

The development of high-throughput sequencing and mass spectrometry technologies, including bulk and single-cell RNA sequencing and quantitative proteomics, has allowed for the systematic measurement of gene and protein expression under different conditions, treatments, and cell types, thereby producing gene lists for further analysis (Haque et al., 2017). Enrichment analysis is a statistical method used to interpret gene lists by examining whether predefined gene sets grouped by a biological pathway or other functional annotations are overrepresented in the observed list beyond what would be expected by chance (Khatri et al., 2012). Databases of prior knowledge such as Gene Ontology (Ashburner et al., 2000) and KEGG (Kanehisa, 2000) among others, contain curated pathways that are widely used by researchers to identify which of their pathways and processes are significant.

Several tools have been developed for this analysis, including DAVID (Huang et al., 2009), Enrichr (Kuleshov et al., 2016), and clusterProfiler (Wu et al., 2021). Although these tools are popular in the community, they only return a list of enriched terms without further detail and ignore the biological questions being asked. Therefore, integrating the biological context to identify the most relevant terms is an additional step that requires a manual search of different databases, literature, or gene functional information.

Large Language Models (LLMs) have been successfully applied in biomedical research for reasoning in complex tasks (Thirunavukarasu et al., 2023; Zhang et al., 2025), with state-of-the-art AI agents combining LLMs with external tools while maintaining a human-in-the-loop (HITL) approach (Qi et al., 2026). It has been shown that LLMs can identify biological functions from gene sets with good accuracy, and their accuracy can be further enhanced by factors such as prior knowledge, tool integration and prompting (Hu et al., 2025). GeneAgent is one such agent (Wang et al., 2025) that uses LLM to annotate gene set functions while using domain databases, including enrichment tools, as a self-verification mechanism to reduce hallucinations. On the other hand, BRAD introduced a framework that includes enrichment analysis as part of an automated biomarker discovery pipeline (Pickard et al., 2025). However, a complete workflow that combines statistical enrichment analysis with biological interpretation while enabling researchers to iteratively guide the analysis has not been developed yet.

Here we present Enrich.AI, the first AI agent platform designed specifically for enrichment analysis, which offers a full workflow from gene sets to the interpretation and visualization of results. The platform can perform enrichment analysis using more than 220 Enrichr libraries, including tools for gene functional annotation, biomedical literature search, and database retrieval. The architecture with the LLM and the tools provided is managed by a reasoning engine built on the LangGraph ReAct agent framework with Google’s Gemini model (Comanici et al., 2025). The model dynamically determines the most appropriate tool combination and interpretation for a specific user query, providing a complete HITL platform where the user can further refine the analysis. The reliability of these interpretations was assessed by evaluating their faithfulness to retrieved evidence and hallucination rate against curated databases, with results showing strong grounding and no fabricated citations.

## 2. Methods

### 2.1. System Architecture

Enrich.AI is based on the ReAct agent framework, which involves a Thought-Action-Observation loop (Figure 1). It employs Google’s Gemini 2.5 Flash as the base model. This model was selected owing to its combination of low-cost efficiency reasoning capabilities, enabling broad accessibility and extended interactions. It is set with a temperature of 0.4 to remain close to the predefined prompt while retaining some freedom to select the optimal workflow.

**Figure 1.**
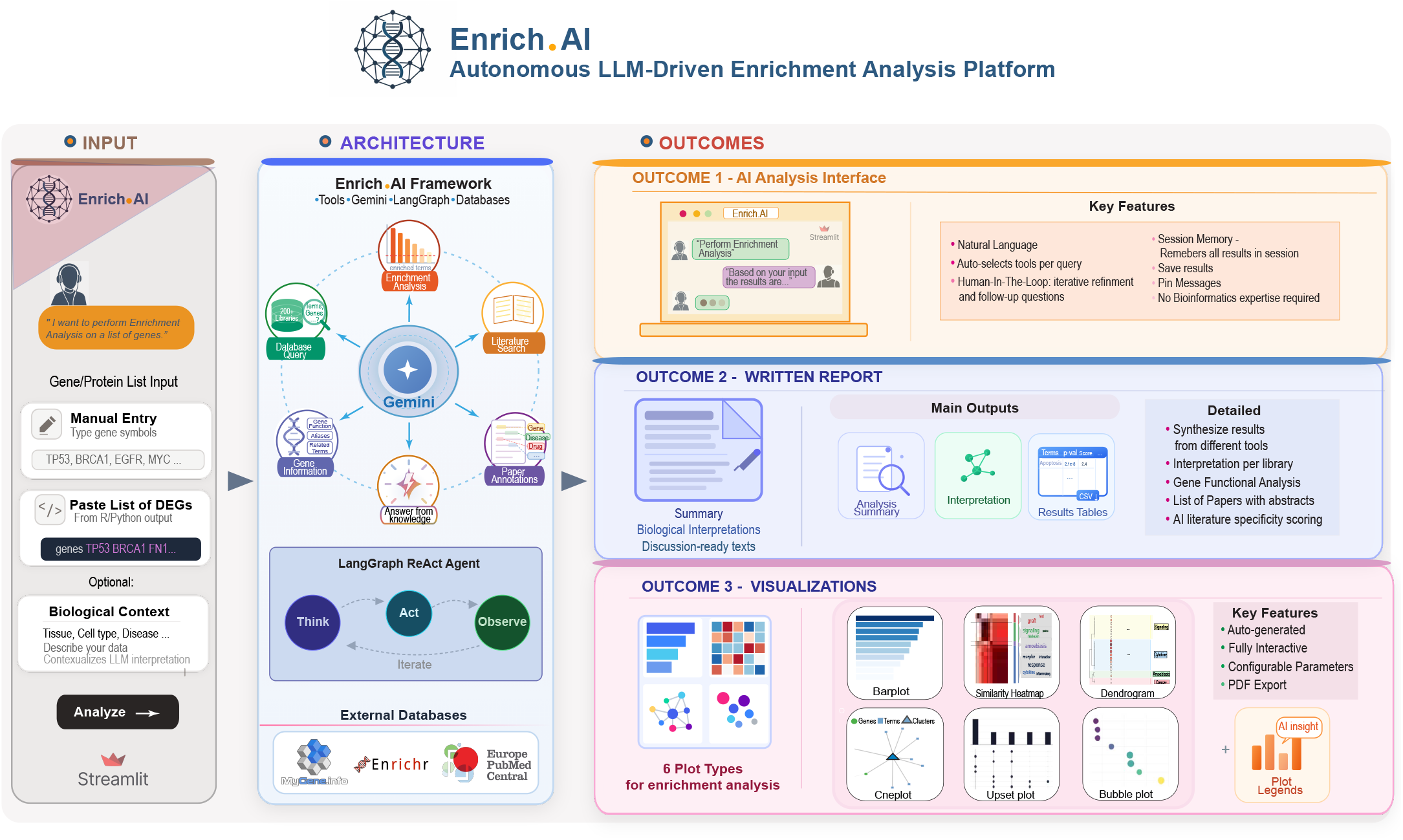
Schematic workflow of Enrich.AI. Input (left): the chat-based user interface, where the researcher enters a gene list and provides optional biological context. Architecture (center): the reasoning engine at the core of the platform. A Gemini 2.5 Flash agent, orchestrated by LangGraph in a ReAct loop, autonomously selects among five biological tools which in turn query external resources. Outcomes (right): Outcome 1 is the interface that supports follow-up questions and human-in-the-loop refinement. Outcome 2 is a written report that synthesizes findings across tools, including biological interpretation. Outcome 3 is a suite of six interactive visualizations for enrichment results.

LangGraph is the orchestrator of the system and connects Gemini to external tools. Following the ReAct workflow, Gemini first reasons about the user prompt and the necessary information required to answer the query, and then selects which tools to use and with what parameters. LangGraph executes the function calls of the tools provided by Gemini and returns the results to the model for observation. The Thought-Action-Observation loop repeats for up to seven hops or until Gemini has gathered all the required information needed, enabling this way the ability for multi-step analyses where multiple tools have to be used. Gemini has access to five tools, and can select one or more based on the user prompt, each registered with a typed Pydantic input schema, which enables valid function calls and reduces any parameter hallucinations. Moreover, the user has access to the execution details and workflow followed by the model, and can verify which tools were invoked and why.

Finally, to provide a full HITL workflow, the model has memory of recent results within a session, enabling iterative refinement of the analysis. It can be used either as a StreamLit web application or run locally.

### 2.2. Enrichment Analysis

For enrichment analysis, Gemini selects the input genes and passes them through LangGraph to Enrichr, where the gene list is tested against user-specified or default libraries. The complete list of over 220 Enrichr libraries is accessible in a separate tab, categorized by disease, phenotype, cell-type signatures, drug perturbations, and genes targeted by transcription factors. Users can either select libraries from this tab or specify them in the prompt. The model uses Enrichr for enrichment analysis, which calculates enrichment using Fisher’s exact test with Benjamini–Hochberg correction, and results are filtered by adjusted *p*-value.

To visualize the results, Enrich.AI provides six plots, several of which were adapted from R packages including clusterProfiler and simplifyEnrichment (Gu and Hübschmann, 2023). The plots are all interactive with a variety of parameters to adjust and fine-tune to produce high-quality visualizations.

### 2.3. Database Retrieval

The database retrieval tool provides multiple functions to access the full collection of 200+ libraries from Enrichr and finds all information requested by the user, such as specific terms, similar terms, and gene-to-term mapping. This tool can be applied when no statistical analysis is required, and Gemini selects the appropriate function to call based on the user query.

### 2.4. Literature Search and Relevance Assessment

Literature search is conducted using the Europe PMC API (The Europe PMC Consortium, 2015), which covers more than 40 million biomedical papers. The Europe PMC returns papers sorted by relevance, publication year, or citation count. Gemini was also given the ability to further evaluate the returned papers by taking into account the title and abstract, producing a specificity score of how relevant they are, with a brief justification. An additional function can focus on full-text papers and retrieve text-mined annotations for genes, diseases, and chemicals.

### 2.5. Gene Information Retrieval

Gene annotations are obtained from the MyGene.info API (Wu et al., 2013), and include the gene name, functionality, aliases, Gene Ontology terms, and KEGG, Reactome, and WikiPathways.

### 2.6. User Interface and Implementation

Enrich.AI is implemented as a web application using the Streamlit framework, providing a chat interface where the user submits queries and receives results. Each response includes a summary, a detailed analysis, and typically a results table, with six visualizations appearing for enrichment analyses. Each response concludes with a reasoning trace that provides full transparency to the agent’s decision workflow. The platform is available with free default access and additionally supports user-supplied API keys across multiple model providers. The application does not collect or share information beyond an active session.

### 2.7. Evaluation

To evaluate the interpretations of the model, the 50 MSigDB Hallmark gene sets were used (Liberzon et al., 2015), with the model performing enrichment analysis and literature search on each. The interpretations were decomposed into claims, and Claude Sonnet 4.5 (Anthropic, 2025) was used as a judge to assess faithfulness to the retrieved evidence, similar to the RAGAS (Es and et al., 2024) framework, hallucination rate on a subset of claims against external databases the tool had no access to, and reproducibility across repeated runs. Two-thirds of claims were grounded in retrieved evidence (65.6%), with the remainder reflecting established background biology (32.1%) and only 2.3% unsupported. The hallucination rate was 2.64% (54/2,047 database-decidable claims), with no fabricated citations across all 90 runs. Enrichment results were consistent across replicate runs, with variation being mainly in the literature search. Further details are provided in the Supplementary Material.

## 3. Discussion

Enrich.AI is a conversational biology assistant made for enrichment analysis and biological interpretation. Compared to existing tools (Table 1), Enrich.AI bridges the gap between existing enrichment analysis tools, which provide lists of terms, and LLMs, which are not statistically rigorous. While this agentic approach is similar to tools such as GeneAgent and BRAD, it tackles a different problem: GeneAgent provides self-verification of LLM predictions for gene annotations, whereas BRAD’s enrichment workflow is an automated pipeline rather than an interactive interpretive session.

**Table 1.**
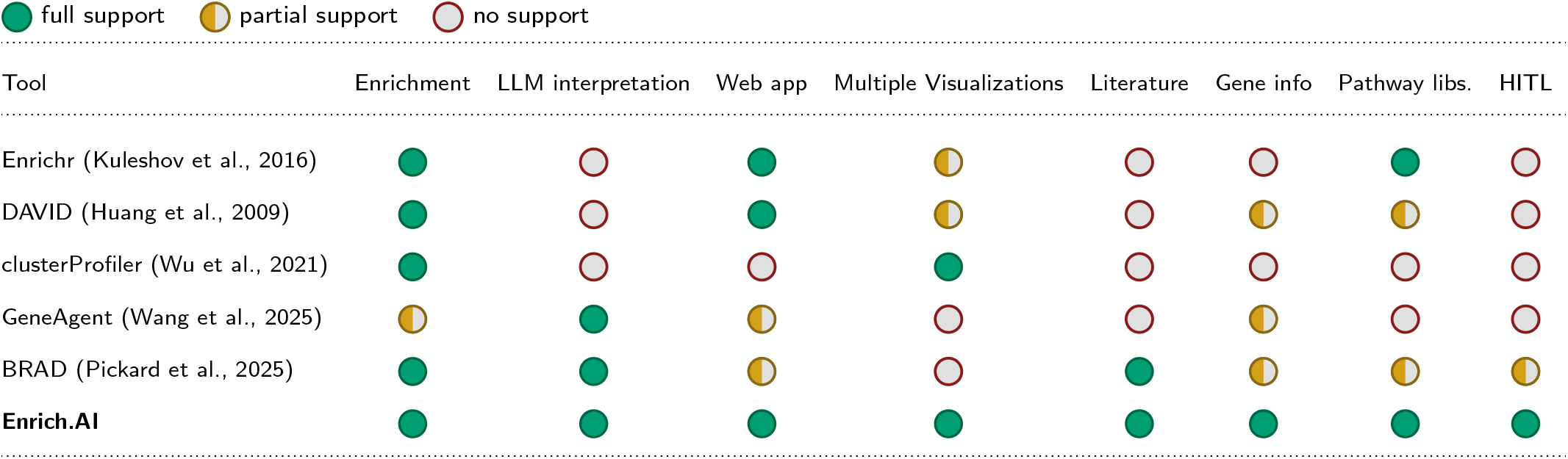
Feature comparison of Enrich. AI with existing enrichment analysis tools and LLM-based agents for gene set analysis.

In addition to interpretation, the system offers six interactive visualizations and access to the entire Enrichr library catalogue, allowing researchers to explore results beyond the default output. The chat interface allows the user to retain control of the reasoning process, which is crucial for a task where the model cannot always make an independent judgment of the biological plausibility of the results.

The platform provides free default access and also accepts user-supplied keys from several model providers. Enrichment is currently performed exclusively through Enrichr, and rank-based approaches such as GSEA have not yet been supported. Future development will focus on incorporating additional enrichment methods and visualizations.

## Notes

https://github.com/jkouprey/Enrich.AI

### Competing Interest Statement

The authors have declared no competing interest.

