## Supplementary Material for "Enrich.AI: An Agentic AI Platform for Interactive Gene Set Enrichment Analysis and Biological Interpretation"

Dimitrios E. Koumadorakis[1,*], Themis P. Exarchos[1], Panagiotis Vlamos[1], Aristidis G. Vrahatis[1]

[1]Bioinformatics and Human Electrophysiology Laboratory, Department of Informatics, Ionian University, 49084, Corfu, Greece

### S1 System Architecture

This section describes the implementation details of the Enrich.AI reasoning engine, including the language model configuration (S1.1), session memory (S1.2), tool registration (S1.3), error handling (S1.4), and system prompt (S1.5).

#### S1.1 Language model

Gemini 2.5 Flash (Comanici et al., 2025) is a sparse mixture-of-experts transformer with a one-million-token context window. The model is configured with a temperature of 0.4 to favor deterministic and reproducible tool selection while allowing flexibility for natural language interpretation, a maximum output of 4,096 tokens for tool-calling steps and 8,192 tokens for the final response synthesis, top-$p$ of 0.95, and top-$k$ of 40. An average single-tool query can take between two and three API calls, whereas a more complex multi-step analysis, which combines enrichment, literature search, and gene information, can take between five and seven calls. The tool is free to use with Gemini 2.5 Flash for limited usage, with the user also having the ability to use their own API key for Gemini 2.5 and 3.5 Flash, Groq's gpt-oss-120b only (Agarwal et al., 2025), as Llama did not give good results, and gpt-5.4-mini from OpenAI. For Groq, the free tier cannot be used, as its output token limit is smaller than the interpretations the tool typically produces, causing responses to be truncated. Although all these models showed good reasoning and tool-calling in our testing, we recommend using Gemini, as it is the model we evaluated and the one we provide freely.

#### S1.2 Conversation memory

The reasoning engine has a memory buffer, which holds the full, uncut output of past tool calls. In the event that the user asks a follow-up query, the context of the last three queries is stored and pre-formatted and prepended to the new query, enabling the model to use the previous results without re-running the analysis. The buffer can hold as many as ten result envelopes.

#### S1.3 Tool registration and input schemas

Each of the five tools is registered with LangGraph as a structured function with a typed Pydantic input schema that declares the name, type, default, and description of every argument. During inference, Gemini produces tool calls in structured JSON that follows these schemas, and the orchestrator validates the arguments and then executes them. Tool calls that have missing required fields, wrong types, or invalid values are rejected prior to reaching the external API and the error of validation is sent back to the model as an observation to enable it to update the call on the subsequent reasoning step. This is a complementary mechanism to the reasoning trace outlined in the main text, as a protection against parameter hallucinations. The full schemas used in the current implementation are shown in Figure S1. Another runtime validation that is unique to the enrichment and database retrieval tools is a comparison of the library names specified by the user or model with the live Enrichr (Kuleshov et al., 2016) library catalog, where a library name that is not found is not silently failed but a structured message listing the possible valid alternatives is returned instead.

#### S1.4 ReAct execution loop and error handling

LangGraph orchestrates the Thought-Action-Observation cycle by routing messages between the language model and the tool executor until either the model produces a final response or the recursion limit of 60 is reached. In case of failures, a structured error message is sent to the model and the agent will correct itself within the same query by re-executing with tweaked parameters or by choosing a different tool. Network errors can be encountered when calling external APIs and are handled at a lower level through an automatic retry mechanism with exponential backoff, configured for up to five attempts on HTTP status codes 429, 500, 502, 503, and 504. A deterministic guard also rejects a final answer that contains no tool call when the query explicitly requested an analysis, prompting the agent to invoke the required tool. The system further contains a fallback path that is initiated at startup, in case the LangGraph agent cannot be built, e.g., because of a failed dependency where in that case, Gemini is invoked directly with the same system prompt and tool descriptions, which can be seen from the execution details or, if run via terminal, in the debug logs. The overall query flow, including these validations, retry, and fallback paths, is shown in Figure S2.

Input-type validation (S1.3), step-by-step error recovery, a deterministic citation guard, and the user-visible reasoning trace offer four complementary countermeasures to reduce and catch the hallucinations that afflict LLM-based analysis tools. Schema validation ensures that invalid tool arguments do not reach the external APIs, error recovery incorporates the ability to repair course within a single query, the citation guard deterministically removes, after generation, any citation not backed by a paper actually retrieved through the literature calling function, so no reference can be fabricated and finally, the reasoning trace makes human verification of each tool call and its outcome possible via he execution details that exist at the end of each tool output. Moreover, the specific temperature used was applied after multiple tests to reduce hallucinations and produce more deterministic results and workflows.

#### S1.5 System prompt

A system prompt is included in the reasoning engine and defines the purpose of the agent and its main principles. The full prompt is reproduced in Figure S3 and is also available in the project repository. The prompt does not tell the model which tool to apply to which question, nor does it dictate a fixed workflow. The model itself reasons and finds the analytical strategy in response to the biological question and does not rely on hard-coded commands.

```
// Tool 1: run_enrichment_analysis (required: genes)
```

```
{
  "type": "object",
  "properties": {
    "genes": { "type": "string",
      "description": "Comma-separated gene symbols (minimum 4 genes)" },
    "libraries": { "type": ["string", "null"], "default": null,
      "description": "Comma-separated Enrichr libraries; null uses defaults" },
    "p_value_threshold": { "type": "number", "default": 0.05,
      "description": "Adjusted p-value cutoff for significance" },
    "top_n": { "type": ["integer", "null"], "default": null,
      "description": "Max terms per library; null returns all significant" }
  },
  "required": ["genes"]
}
```

```
// Tool 2: get_gene_info (required: gene)
```

```
{
  "type": "object",
  "properties": {
    "gene": { "type": "string",
      "description": "Gene symbol (e.g., 'TP53', 'BRCA1', 'EGFR')" },
    "organism": { "type": "string", "default": "9606",
      "description": "NCBI taxonomy ID; default 9606 (human)" }
  },
  "required": ["gene"]
}
```

```
// Tool 3: db_retrieve (required: query)
```

```
{
  "type": "object",
  "properties": {
    "query": { "type": "string",
      "description": "Search query: symbols, term names, IDs, or keywords" },
    "libraries": { "type": ["string", "null"], "default": null,
      "description": "Comma-separated Enrichr library names" },
    "task": { "type": "string", "default": "auto",
      "enum": ["auto", "term_search", "term_details", "term_genes",
               "gene_terms", "library_browse", "find_similar"],
      "description": "Query mode; auto lets the agent choose" },
    "organism": { "type": "string", "default": "9606",
      "description": "NCBI taxonomy ID" },
    "limit": { "type": "integer", "default": 20,
      "description": "Maximum results to return" },
    "include_genes": { "type": "boolean", "default": false,
      "description": "Whether to include gene lists per term" }
  },
  "required": ["query"]
}
```

```
// Tool 4: search_literature (required: query)
```

```
{
  "type": "object",
  "properties": {
    "query": { "type": "string",
      "description": "Europe PMC search query" },
    "max_results": { "type": "integer", "default": 20,
      "description": "Maximum papers to return (up to 100)" },
    "min_year": { "type": ["integer", "null"], "default": null,
      "description": "Minimum publication year filter" },
    "max_year": { "type": ["integer", "null"], "default": null,
      "description": "Maximum publication year filter" },
    "sort_by": { "type": "string", "default": "relevance",
      "enum": ["relevance", "citations", "date"],
      "description": "Sort order for results" }
  },
  "required": ["query"]
}
```

```
// Tool 5: get_paper_annotations (required: paper_id)
```

```
{
  "type": "object",
  "properties": {
    "paper_id": { "type": "string",
      "description": "Paper identifier: PMID, PMCID, or DOI" },
    "id_type": { "type": "string", "default": "auto",
      "enum": ["pmid", "pmcid", "doi", "auto"],
      "description": "Identifier type; auto detects automatically" },
    "annotation_types": { "type": ["string", "null"], "default": null,
      "description": "Comma-separated annotation types; null returns all" }
  },
  "required": ["paper_id"]
}
```

**Figure S1.** JSON schemas for the five tools registered with the reasoning agent. Each schema is derived from a Pydantic model and validated against Gemini's tool calls before execution.

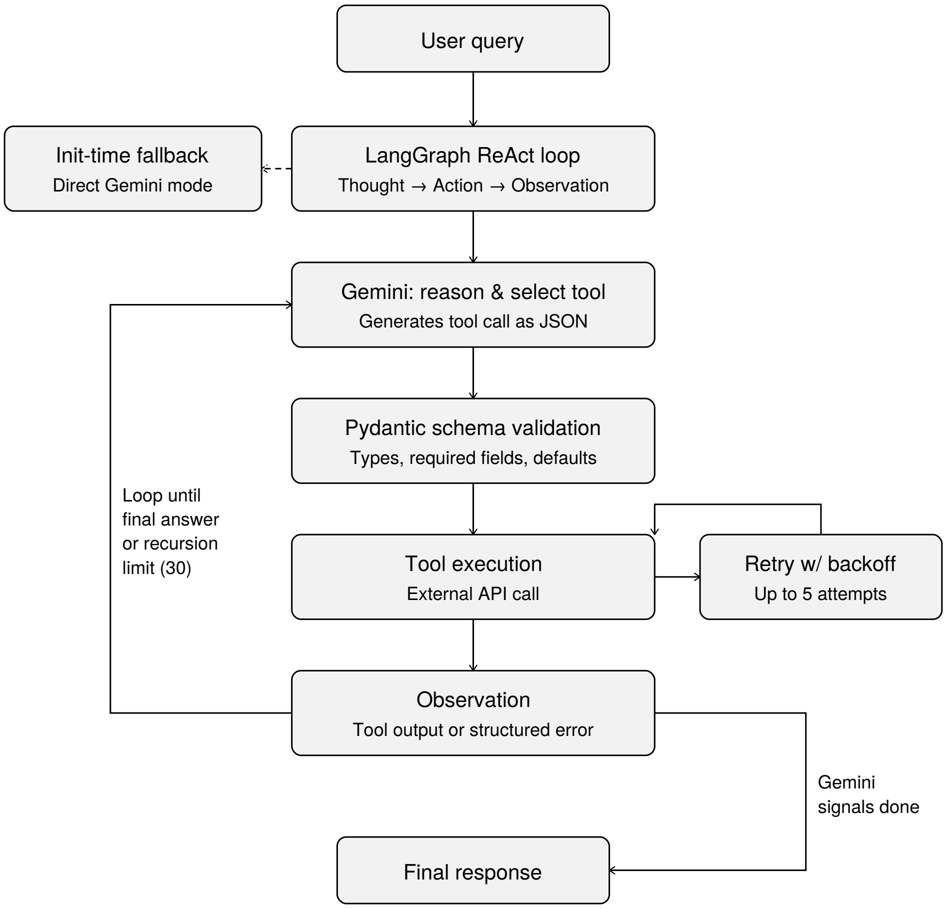


**Figure S2.** Query execution flow in Enrich.AI. A user query enters the LangGraph ReAct loop, where Gemini reasons about the biological question and generates a tool call as structured JSON. Pydantic schema validation intercepts calls with invalid arguments before execution. The selected tool calls an external API; transient HTTP errors trigger automatic retry with exponential backoff (up to five attempts on status codes 429, 500, 502, 503, 504). The tool's output, or a structured error message on failure, is returned as the observation, and the loop continues until Gemini produces a final response or the recursion limit is reached. An initialization-time fallback path (dashed arrow) activates if the LangGraph agent cannot be constructed, in which case Gemini is invoked directly with the same tool descriptions.

```
// Enrich.AI system prompt

You are Enrich.AI, an expert biology research assistant for molecular biologists. You have deep
     knowledge of genomics, signaling pathways, disease mechanisms, systems biology, and
     experimental design.

You have access to tools for enrichment analysis, gene information, literature search, database
     queries, and paper annotation. You may call any tool, in any order, as many times as
     needed. You decide when and why.

=== CORE PRINCIPLE ===

Enrichment tools return lists. You provide biological meaning.

Your role is to interpret what those pathways mean in the specific experimental context,
     identifying what is expected, mechanistically coherent, surprising, potentially
     artifactual, and worth follow-up.
When biological context is provided (e.g., tissue, disease, condition), your primary task is to
     prioritize: rank the enriched terms by biological relevance to that context, explain why
     certain terms matter more than others, and flag which are noise or generic.
You have to be detailed in your answer, highlighting terms based on context is your main aim and
     you have a variety if tools to validate your answer.
You are a knowledgeable colleague reviewing results together and sharing your experience.

=== HOW YOU THINK ===

You reason, then act, then observe, then reason again. There is no fixed pipeline.

- Think about what biological question is being asked.
- Decide which tools are appropriate.
- After results return, interpret before acting again.
- If something looks incomplete, inconsistent, or intriguing -- investigate further.
- If a tool fails or returns weak signal, adapt.
- Let the biology determine your workflow.

=== VALIDATE WHAT YOU CLAIM IF NEEDED ===

Do not state biological conclusions without evidence. Use your tools to support your
     interpretations:
- Use search_literature to find papers that confirm or challenge a mechanistic link.
- Use db_retrieve with short keyword queries to check pathway membership or term overlap.
- Use get_gene_info to verify gene functions before making claims about their role.
- If a paper has full text available, you can retrieve it for deeper validation.
Your credibility depends on grounding interpretation in data, not generating plausible-sounding
     text.

=== FOLLOW-UP QUESTIONS ===

Look your memory to see what you have. Does the user ask about previous results?
If yes, do NOT re-query tools to find information you already have. Use tools only to validate
     or expand,
not to re-discover what is already in the conversation context.

=== CONTEXT IS EVERYTHING ===

The same enriched term can mean different things depending on the tissue, disease, perturbation,
     species, or experimental design. Always anchor interpretation to the biological context.
If context is not explicitly provided, infer it from the gene signature itself, read it like a
     biologist would.

=== EVIDENCE AND CONFIDENCE ===

Distinguish clearly between strong conclusions supported by data, plausible mechanistic
     hypotheses, and speculative interpretations. If you use literature to answer cite the
     papers. Maintain scientific credibility.

=== RESPONSE GUIDELINES ===

- Structure your response with a summary section first, then use "## Detailed Analysis" as a
     header
  before the in-depth interpretation. The summary and detailed sections will be displayed
     separately in the UI.
- Structure responses based on what the query needs. Use headers.
- Do not output raw markdown tables.
- For db_retrieve use short keyword queries, not full sentences.
- State which tools were used and what parameters were applied.
- Discuss results from ALL tools called.
- If the first approach fails, try at least one alternative before reporting failure.
- For get_gene_info, be selective. Do not look up only one or every gene in a list if you have
     hundreds of them. Focus on the most
  biologically informative or ambiguous genes that would change interpretation.
```

**Figure S3.** System prompt initializing the reasoning engine. The prompt is intentionally concise and behavioral rather than prescriptive: it establishes the agent's role and core principle but does not specify a tool-selection pipeline, leaving workflow decisions to the model at inference time.

### S2 Enrichment Analysis Details

Enrichr computes enrichment using Fisher's exact test, which evaluates whether the overlap between the user's gene list and a predefined gene set, such as a biological pathway, is greater than expected by chance. For a given term, let $k$ denote the number of input genes that overlap with the term's gene set, $K$ the total number of genes annotated to that term in the library, $n$ the number of genes in the user's input list, and $N$ the total number of genes in the background. Fisher's exact test computes the probability $p$ of observing an overlap of $k$ or more genes under the null hypothesis of random gene selection. Multiple testing correction is applied across all terms within each library using the Benjamini-Hochberg method, producing an adjusted $p$-value for each term. Terms with an adjusted $p$-value above the threshold (default 0.05) are excluded from the results.

For each term that passes the significance threshold, Enrichr also computes a $z$-score $z$, which represents the deviation of the term's rank from its expected rank across permutations of random gene sets. A large negative $z$-score indicates that the term ranks considerably higher than expected by chance. The combined score integrates both metrics and is defined as:

$$c = \log(p) \cdot z \tag{S1}$$

where $p$ is the raw Fisher's exact test $p$-value and $z$ is the $z$-score. The product yields a positive score where higher values indicate stronger enrichment.

The overlap for each term is expressed as $k/K$, where $k$ and $K$ are defined as above, providing a measure of how much of the pathway is covered by the input gene list. The list of overlapping genes is also returned, enabling downstream inspection of which specific input genes drive the enrichment of each term. Results are sorted by adjusted $p$-value in ascending order, with the most significant terms appearing first.

### S3 Database Retrieval

The database retrieval tool provides access to all 200+ libraries taken from the Enrichr API. It supports six modes that Gemini selects automatically based on the user query. Term search retrieves terms matching a keyword across one or more libraries, term details return full information for a specific term including its gene set, gene-to-term mapping retrieves all terms containing a given gene, and term-to-gene mapping returns all genes annotated to a given term. Moreover, library browsing lists all available terms within a specified library, and similarity search finds terms whose gene sets are most similar to a query term using Jaccard similarity.

### S4 Visualization Details

Enrich.AI provides six interactive visualization types, several of which were in R, including clusterProfiler (Wu et al., 2021) and simplifyEnrichment (Gu and Hübschmann, 2023) and implemented in Python. All plots are fully interactive with zooming, panning, and hover tooltips. Apart from interactive plots, a variaty of parameters is also included to tune such as terms to select, color, x axis and width and height ratios. Each plot has an AI interpretation button that sends the current plot to Gemini to provide a legend. All plots can be exported in either PDF or PNG format.

#### S4.1 Bar plot and bubble chart

The bar plot shows enriched terms as horizontal bars ranked by a user-selected metric such as gene count, $-\log_{10}$ (adjusted $p$-value), or combined score, and color can be one of the above parameters or libraries. The bubble chart extends this representation by encoding an additional dimension through bubble size, which is proportional to gene count. In both visualizations the user can select which terms to include and the color palette to use. Example outputs are shown in Figures S4 and S5.

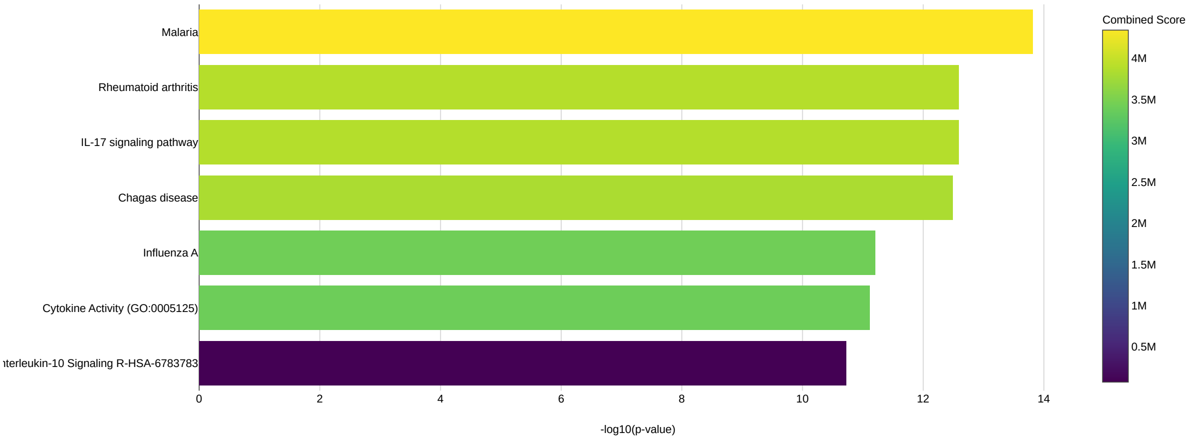


**Figure S4.** Example bar plot output from Enrich.AI. Enriched terms are displayed as horizontal bars, with bar length proportional to the $-\log_{10}$ (adjusted $p$-value) and color mapped to the combined score.

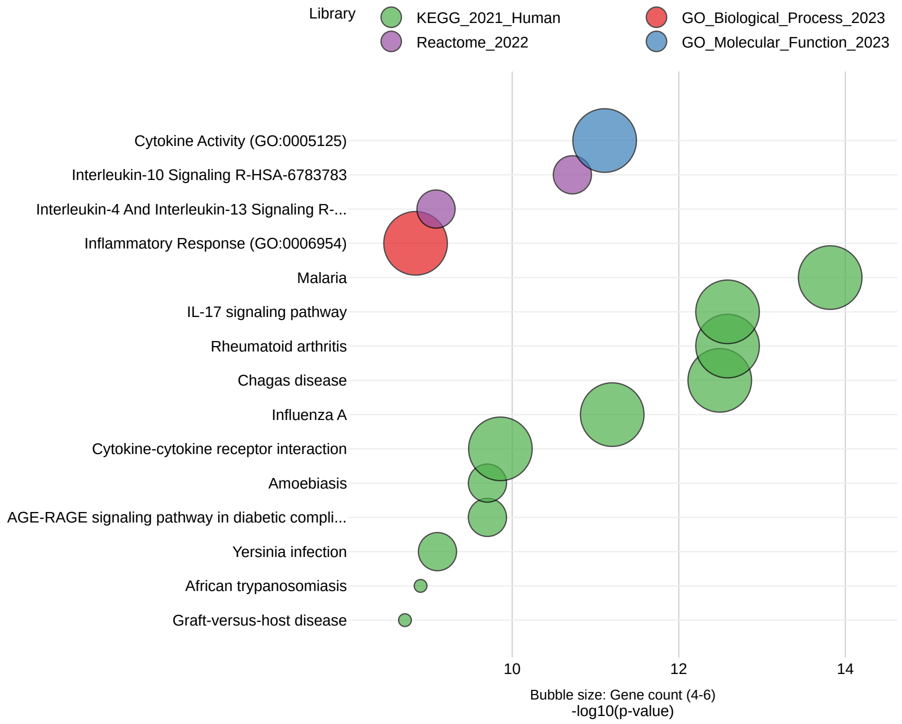


**Figure S5.** Example bubble chart output from Enrich.AI. Each enriched term is represented by a bubble positioned along the $-\log_{10}$ (adjusted $p$-value) axis, with bubble size proportional to gene count and color indicating the source library.

### S4.2 UpSet plot

The UpSet plot is a way to view intersection patterns of gene sets across selected terms. For each gene in the union of the terms, a binary vector is generated that indicates the terms in which the gene is found. The plot is divided into two panels, where the top panel displays the size of the intersections as a bar chart, with each bar representing the number of genes shared among these sets, and the bottom panel displays the membership patterns of the intersections as a dot matrix with filled dots denoting which terms are involved in each intersection. An example is shown in Figure S6.

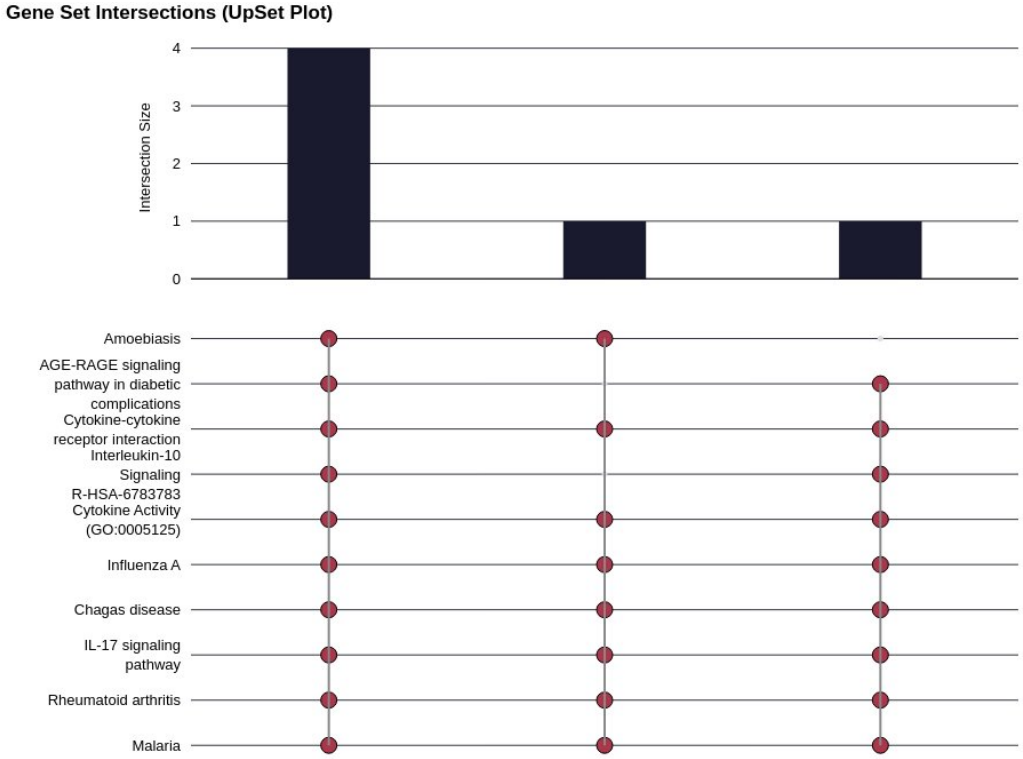


**Figure S6.** Example UpSet plot output from Enrich.AI. The upper panel shows intersection sizes; the lower panel displays the membership pattern for each intersection.

### S4.3 Common computational foundation

The methodological approach used in the rest of the visualizations, specifically the dendrogram, the similarity heatmap, and the concept network plot, is based on the same calculation. For each pair of enriched terms, their similarity is computed using the Jaccard index over their annotated gene sets:

$$J(A,B) = \frac{|A \cap B|}{|A \cup B|} \tag{S2}$$

where $A$ is the set of overlapping genes for the first term and $B$ is the set of overlapping genes for the second term. A Jaccard index of 1 means that the two gene sets are the same and 0 means that they do not overlap. The similarity matrix is transformed into a distance matrix $D = 1 - J$ and then agglomerative hierarchical clustering is applied using Ward's minimum variance approach (Ward Jr, 1963) as implemented in SciPy (Virtanen et al., 2020). Ward's method starts from each term and works by minimizing the total within-cluster variance at each step by merging the clusters that produce the smallest increase in variance. The number of clusters is determined using the maxclust criterion, targeting approximately five terms per cluster by default. The Cluster names are generated automatically by tokenizing the term names within each cluster, removing generic words, and selecting the most frequent remaining keywords by count.

### S4.4 Hierarchical clustering dendrogram

The dendrogram displays hierarchical clustering of enriched terms using the Jaccard gene set similarity, similar to clusterProfiler's dendrogram. The top $n$ most significant terms (ranked by adjusted $p$-value) are chosen and clustered as described in S4.3 with a user-specified linkage method (Ward, complete, or average). The figure layout consists of five elements from left to right. On the left, the tree shows the hierarchical relationships between terms, with shorter branches indicating higher gene set similarity between the connected terms. The background rectangles show the clusters, each with a unique color. The leaves of the tree are the terms themselves, depicted as dots whose color intensity reflects their statistical significance and whose size reflects the number of input genes overlapping with that term. Finally, cluster labels are placed at each cluster, automatically generated by extracting keywords from grouped terms to describe their common biological functions. An example is shown in Figure S7.

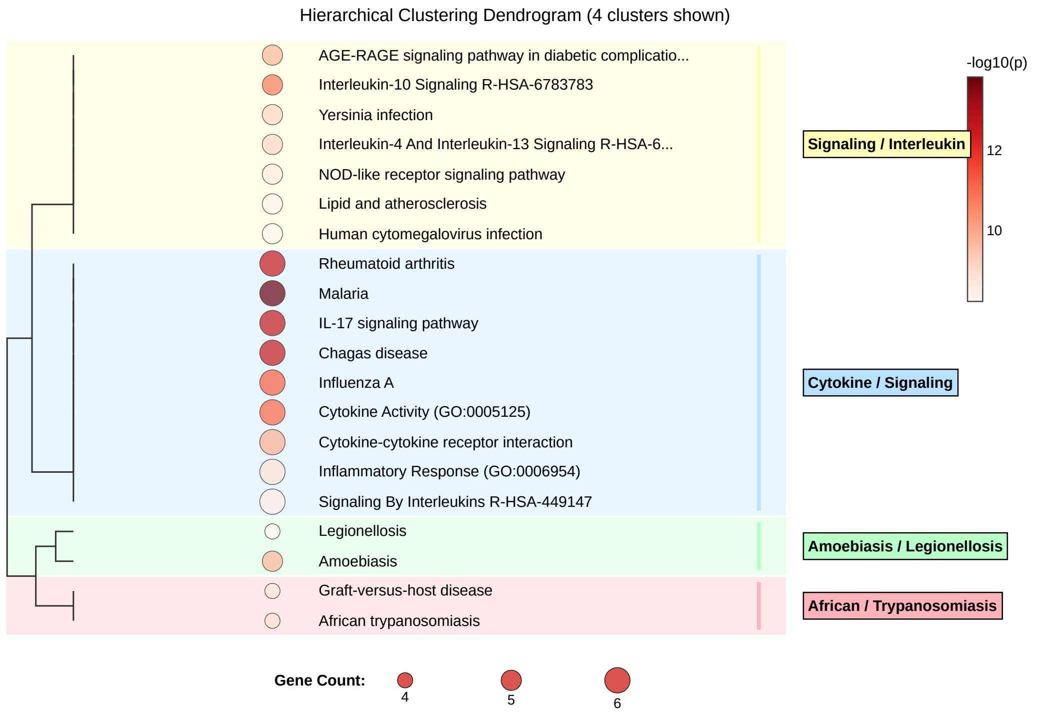


**Figure S7.** Example hierarchical clustering dendrogram output from Enrich.AI. Terms are grouped into clusters based on Jaccard similarity of their gene sets; dot color encodes $-\log_{10}$ (adjusted $p$-value) and dot size encodes gene count. Cluster labels on the right are generated automatically by keyword extraction from the term names within each cluster.

### S4.5 Similarity heatmap with keyword annotation

This visualization was inspired by simplifyEnrichment and displays the pairwise Jaccard similarity matrix in a form of a heatmap whose terms are reordered with hierarchical clustering. The top 10 terms by adjusted $p$-value are selected from each cluster to select the most frequent keywords, which are then displayed in a panel. The font size is proportional to the frequency of the word with more frequent ones having a higher font size. An example visualization is shown in Figure S8.

### S4.6 Concept network plot (cnetplot)

The cneplot as implemented in clusterProfiler and adopted here as well, is a bipartite network linking enriched term clusters to their constituent genes.

Terms are clustered based on Jaccard similarity. The network contains three node types: cluster centers produced by Jaccard similarity represented as triangles labeled with keyword-derived names, individual terms and pathways represented as squares with the same color as their cluster and size proportional

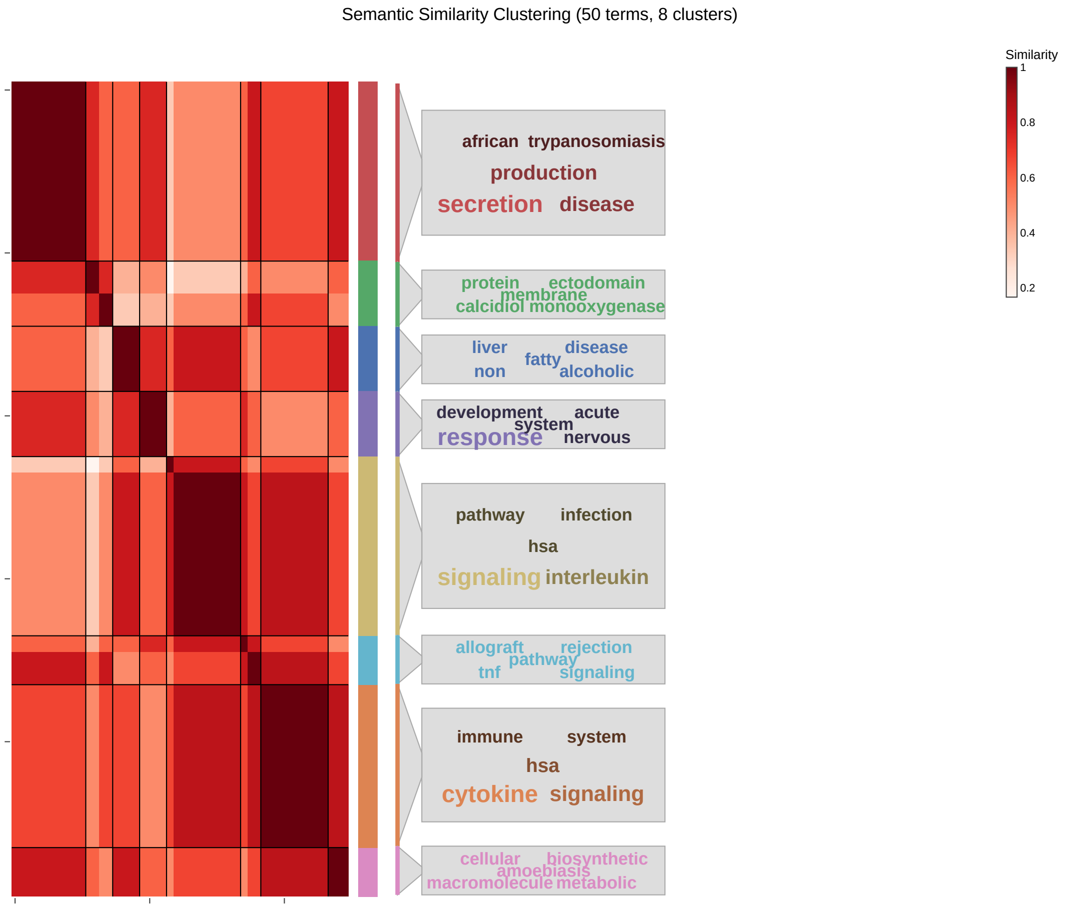


**Figure S8.** Example similarity heatmap output from Enrich.AI. Each cell shows the pairwise Jaccard similarity between two enriched terms; terms are reordered by hierarchical clustering. The colored column on the right indicates cluster membership, and the annotation panels display the top keywords extracted from each cluster's terms.

to their gene count, and genes represented as circles. For the layout, as a first step, the cluster centers are positioned with their terms and genes placed on a radius around them and then the layout is refined using a spring-based layout to minimise edge crossings while retaining the cluster structure.An example is shown in Figure S9.

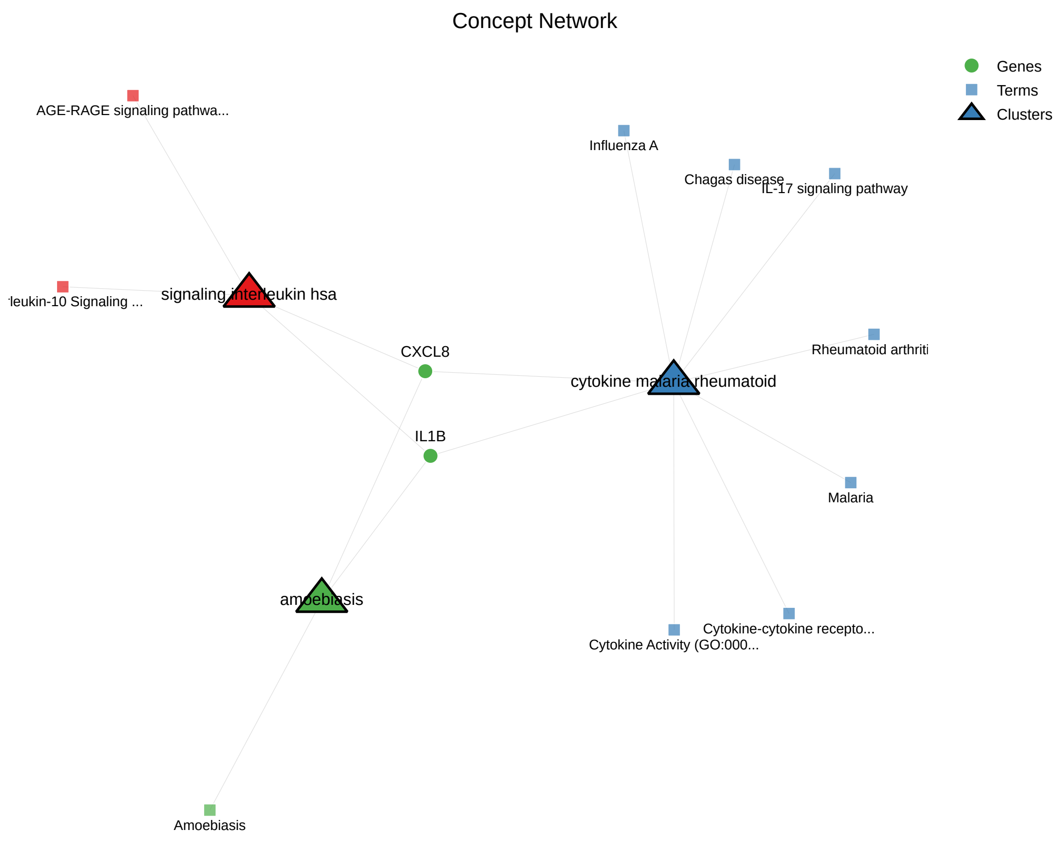


**Figure S9.** Example concept network plot output from Enrich.AI. Cluster centers with keyword-derived names are shown as triangles, individual enriched terms colored by cluster membership are shown as squares, and circles denote genes. Edges connect terms with their cluster center and genes with the clusters that their terms belong to.

### S5 Default Parameters and Configuration

Since it is important in research for workflows to be reproducible, all the default parameters used by the model are summarized here. If no libraries are selected for enrichment analysis five will be used as default, covering the major functional categories:

- `GO_Biological_Process_2023`
- `GO_Molecular_Function_2023`
- `GO_Cellular_Component_2023`
- `KEGG_2021_Human`
- `Reactome_2022`

The adjusted $p$-value significance threshold is 0.05 and all significant terms below that threshold are returned. Literature search returns the top 20 papers from Europe PMC, ranked by relevance. Database retrieval returns up to 20 results per query.

The reasoning loop is bounded by a LangGraph recursion limit of 60 and a memory buffer of 10 past envelopes, with the three most recent queries included as context for each follow-up.

For visualizations, the default number of terms displayed is 25 for the dendrogram and concept network plot, 8 for the UpSet plot, and 50 for the similarity heatmap. The clustering uses Ward's linkage with the maxclust criterion, targeting approximately five terms per cluster, and the concept network plot uses five clusters by default. All of these parameters can be changed per analysis by the user.

### S6 Evaluation

To evaluate the model, we downloaded 50 gene sets of the MSigDB Hallmark collection (Liberzon et al., 2015) and performed three different evaluation checks to understand whether the interpretations it returns are reliable. First, we checked evidence faithfulness, which is whether the interpretation is based on claims the agent retrieved from the tools it used. The second metric addressed factual correctness, where a portion of the model's claims were tested against databases the tool never accessed, to validate whether the claims were correct. The final metric was reproducibility, where we checked the model's evidence faithfulness across repeated runs. For the evaluation we used Gemini 2.5 Flash as the model and Claude Sonnet 4.5 (Anthropic, 2025) as the judge, at zero temperature for reproducibility, to perform claim decomposition and verification of the claims against the databases. Metrics are summarized in Table S1.

For evidence faithfulness, we used an architecture similar to RAGAS (Es et al., 2024). We ran the model on the 50 gene sets, each time asking it to perform enrichment analysis and provide an interpretation supported by literature. The full output, comprising the significant enrichment terms and the retrieved literature with paper abstracts, was then used by Claude Sonnet to decompose these interpretations into individual claims. This yielded a total of 4,737 claims from 45 gene sets, as 5 of these runs did not use any tool. To evaluate the model's faithfulness to the retrieved material, we split the claims into supported claims (grounded and faithful), background claims (drawn from the tool's own knowledge and assessed as correct), and unsupported claims (not backed by the retrieved evidence). From the produced results, the faithfulness of our model was 65.6% [95% CI 64.2-67.0], with background at 32.1% and unsupported claims at 2.3%, showing that more than two-thirds of the claims are grounded in the tools themselves, that the great majority are biologically established, and that only around 2% are unsupported. To validate that this evaluation was consistent, we assessed the judge's reproducibility. Eight gene sets were selected and evaluated a second time, with the judge yielding 98% agreement and confirming that the reported results are consistent.

Since faithfulness captures only the retrieved context, and pathway-level annotations can support claims that are wrong at the level of protein function, we further verified potential hallucinations against the curated UniProt (uni, 2025), NCBI Gene (Brown et al., 2015), and Gene Ontology databases (Ashburner et al., 2000). For this check, we tested the 2,047 claims that named specific genes and pathways. The judge compared each of these claims against the database records and flagged 4.2% [95% CI 3.41-5.16] as potential errors. After revising and adjudicating the flagged claims, 2.64% [95% CI 2.03-3.43] were confirmed as genuine biological errors. To further validate this, a random sample of correct claims was evaluated for false positives, and none were found.

Lastly, we assessed the faithfulness of the model across repeated runs. From the 50 gene sets, 10 were randomly selected and rerun 5 times each. The enrichment results were consistent throughout, using the same libraries and, for standard calls, returning the same terms; the main variation was in the literature search, as the model issued different queries or a different number of calls.

It is also worth noting that across all runs the citation guard operated as intended, and no fabricated citations were found across the 90 runs.

**Table S1.** Evaluation metrics for the Enrich.AI interpretation layer across 50 MSigDB Hallmark gene sets (tool under test: Gemini 2.5 Flash; judge/verifier: Claude Sonnet 4.5). Bracketed values are 95% Wilson confidence intervals.

| Metric | Value | Basis |
|---|---|---|
| *Evidence faithfulness* | | |
| Supported | 65.6% [95% CI 64.2–67.0] | 3,108/4,737 claims |
| Background | 32.1% [95% CI 30.8–33.4] | 1,520/4,737 claims |
| Unsupported | 2.3% [95% CI 1.9–2.8] | 109/4,737 claims |
| *Hallucination rate vs. databases* | | |
| Corrected | 2.64% [95% CI 2.03–3.43] | 54/2,047 claims |
| Original | 4.20% [95% CI 3.41–5.16] | 86/2,047 claims |
| *Other metrics* | | |
| Fabricated citations | 0% | 0/90 runs |
| Tool-less rate | 10.0% | 50-set run |

### S7 HITL Use-Case Scenario

Here we illustrate a simple use-case scenario which combines enrichment analysis and a context-specific follow-up query (Figure S10). A user submits a set of genes and requests an enrichment analysis. Enrich.AI produces the analysis results together with an interpretation of them and six interactive visualizations. In the next step, the user poses a context-specific follow-up: from the KEGG pathways that were found, which of them are related to tumor progression. Instead of re-running the analysis, the model reasons to select the existing results, uses its prior knowledge to narrow the terms to candidate cancer-linked pathways, and validates each of these terms through a literature search before synthesizing the final answer, something that is available for the user to see in the execution details. The interpretation of the model ranks the pathways based on their relevance, and associated with real papers found in the literature, and provides reasoning. In that scenario, IL-17 signaling pathway, TNF signaling pathway and Inflammatory bowel disease, along with retrieved citations (Lin et al., 2026; Qin et al., 2025; Ren et al., 2025; Singh et al., 2026) were ranked in the higher positions.

After Enrich.AI identified the pathways of interest, the user can return to the visualizations that were produced during the first prompt interaction and select the terms of interest to visualize. That way, the user now has six different context-specific visualizations to choose from, along with an AI-generated legend if wanted and a justification to use from the interpretation text of the model. This scenario demonstrates one simple human-in-the-loop design of the platform in which the model autonomously selects the tools to use and in what way while the user steers the analysis toward a specific question of interest.

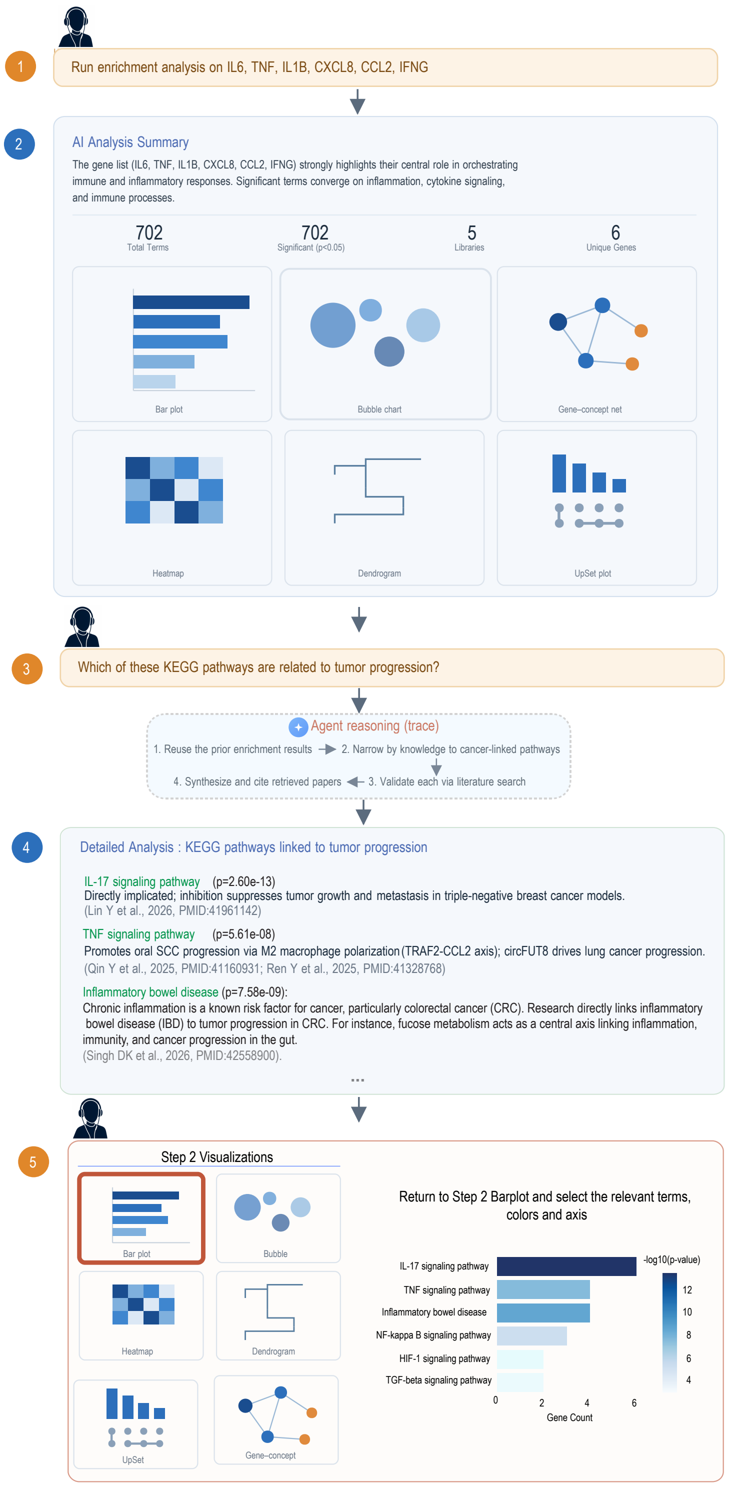


**Figure S10.** Use-case scenario. The user sends a query (step 1) for enrichment analysis, and the tool returns, at step 2, the results with the interpretation and the six visualizations. The user then asks a context-specific follow-up question (step 3), and the tool reasons with the results it has and what it wants to do to answer this question, and returns the interpretations and the tools it called at step 4. The user then returns to the visualizations, selects the bar plot, and selects the relevant terms based on the context, illustrating a simple human-in-the-loop refinement.

### S8 Considerations

Enrich.AI is designed as a biological assistant for enrichment analysis that can interpret the results and provide feedback to the user to identify relevant terms, search literature and having access to more than 200 libraries. Although the system contains various tools to provide accurate feedback, it still relies on a language model whose outputs may still contain inaccuracies. It is important for the user to consult the reasoning trace to support verification of the tools and parameters used, if any.

### Additional Material

A video demonstrating how the platform works is available at: `https://youtu.be/ngJ2fmoXQ6o` and a demo video available at: `https://youtu.be/NBRS4QtYbwg`.
